# DeeplyTough: Learning Structural Comparison of Protein Binding Sites

**DOI:** 10.1101/600304

**Authors:** Martin Simonovsky, Joshua Meyers

## Abstract

**Motivation:** Protein binding site comparison (pocket matching) is of importance in drug discovery. Identification of similar binding sites can help guide efforts for hit finding, understanding polypharmacology and characterization of protein function. The design of pocket matching methods has traditionally involved much intuition, and has employed a broad variety of algorithms and representations of the input protein structures. We regard the high heterogeneity of past work and the recent availability of large-scale benchmarks as an indicator that a data-driven approach may provide a new perspective.

**Results:** We propose DeeplyTough, a convolutional neural network that encodes a three-dimensional representation of protein binding sites into descriptor vectors that may be compared efficiently in an alignment-free manner by computing pairwise Euclidean distances. The network is trained with supervision: (i) to provide similar pockets with similar descriptors, (ii) to separate the descriptors of dissimilar pockets by a minimum margin, and (iii) to achieve robustness to nuisance variations. We evaluate our method using three large-scale benchmark datasets, on which it demonstrates excellent performance for held-out data coming from the training distribution and competitive performance when the trained network is required to generalize to datasets constructed independently.

**Availability:** https://github.com/BenevolentAI/DeeplyTough

## 1 Introduction

Analysis of the three-dimensional (3D) structures of proteins, and in particular the examination of functional binding sites, is of importance in drug discovery and biological chemistry (Skolnick and Fetrow, 2000; Ferreira *et al.*, 2015). Binding site comparison, also known as pocket matching, can be used to predict selectivity of ligand binding, as an approach for hit-finding in early drug discovery or to suggest the function of, as yet, uncharacterized proteins (Ehrt *et al.*, 2016). Structural approaches for pocket matching have been shown to be more predictive of shared ligand binding between two proteins than global structure or sequence similarity (Grishin, 2001; Illergård *et al.*, 2009). Increasingly, there is interest in applying pocket matching approaches to large datasets of protein structures to enable proteome-wide analysis (Holm and Sander, 1996; Hou *et al.*, 2005; Degac *et al.*, 2015; Meyers *et al.*, 2016; Bhagavat *et al.*, 2018). Existing approaches for quantifying the structural similarity between a pair of putative protein binding sites exhibit a range of hand-crafted pocket representations, as well as a combination of alignment-dependent and alignment-free algorithms for comparison (Ehrt *et al.*, 2016; Naderi *et al.*, 2018).

A key measure of success for a pocket matching algorithm is the ability to assign similarity to pairs of protein pockets that have been shown to bind identical ligands (Barelier *et al.*, 2015; Chen *et al.*, 2016). This requirement is useful since it encourages pocket similarity towards biological relevance, however the binding of identical ligands to unrelated pockets is highly dependent on the nature of the ligand (as well as the protein) and often there is no common structural pattern between the pair of binding sites (Meyers *et al.*, 2018; Pottel *et al.*, 2018). As noted by Barelier *et al.* (2015): “The same ligand might be recognized by different residues, with different interaction types, and even different ligand chemotypes may be engaged". It is therefore unsurprising that varied algorithms for pocket matching differ in the manner by which cavities are represented, or as to how different feature types are weighted in the resulting measure of similarity. We argue that a solution that is able to learn from data is expected to perform well since this offers the possibility to remove bias associated with hand-engineered protein pocket representations and their matching, others have also expressed this view (Naderi *et al.*, 2018).

Deep learning has become the standard machine learning framework for solving many problems in computer vision, natural language processing, and other fields (LeCun *et al.*, 2015). This trend has also reached the community of structural biology and computational chemistry, showing utility in a range of scenarios relevant to drug discovery (Rifaioglu *et al.*, 2018). In particular, trainable methods have been applied to protein structure data for a number of applications including protein-ligand affinity prediction (Wallach *et al.*, 2015; Gomes *et al.*, 2017; Ragoza *et al.*, 2017; Stepniewska-Dziubinska *et al.*, 2018; Jiménez *et al.*, 2018; Imrie *et al.*, 2018), protein structure prediction (AlQuraishi, 2018; Evans *et al.*, 2018), binding pocket inpainting (Škalicĉ *et al.*, 2019), binding site detection (Jiménez *et al.*, 2017) and prediction of protein-protein interaction sites (Fout *et al.*, 2017; Townshend *et al.*, 2018). However, to our knowledge a deep learning approach to pocket matching has not been previously described.

A challenge for any machine learning-based method for pocket matching is presented by the available quantity of known protein pocket pairs and the quality of their annotations. Conveniently, Govindaraj and Brylinski (2018) have recently compiled TOUGH-M1, a large-scale dataset of roughly one million pairs of pockets differentiated by whether or not they bind structurally similar ligands. In this paper, we rely on this collection for training and expect that the scale will help the method overcome the noise inherently present in automated strategies for gathering data.

We introduce DeeplyTough, a pocket matching method that uses a convolutional neural network (CNN) to encode protein binding sites into descriptor vectors. Once computed, descriptors can be compared very efficiently in an alignment-free way by simply measuring their pairwise Euclidean distances. This efficiency makes the proposed approach especially suited to investigations on large datasets. Our main contribution is the formulation of pocket matching as a supervised learning problem with three goals: i) to provide similar pockets with similar descriptors, ii) to separate the descriptors of dissimilar pockets by a minimum margin, and iii) to achieve robustness to selected kinds of nuisance variability, such as specific delineations of binding sites. We thoroughly evaluate our method on three recent large-scale benchmarks for pocket matching. Concretely, we demonstrate excellent performance on held-out data coming from the training distribution (TOUGH-M1) and competitive performance when the trained network is required to generalize to datasets constructed independently by Chen *et al.* (2016) and Ehrt *et al.* (2018).

## 2 System and Methods

The problem of protein pocket matching is seen from the perspective of computer vision in our approach. The main idea is to regard pockets as 3D images and process them using a CNN to obtain their representation in a vector space where proximity indicates structural similarity, see Figure 1. In this section, we first discuss the choice of the training dataset and featurization. Then, we pose our method as a descriptor learning problem and describe a training strategy that encourages robustness to nuisance variability. Finally, we describe the architecture of the neural network and details relevant to its implementation.

**Fig. 1.**
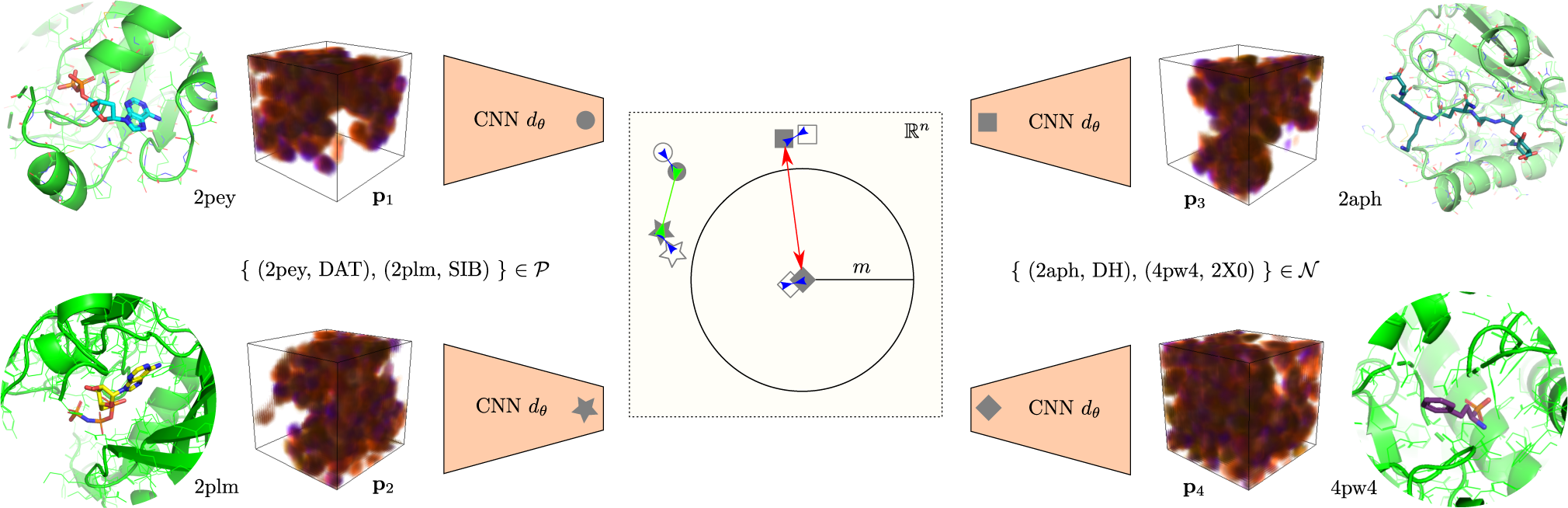
Illustration of learning binding site matching with a contrastive loss function (green and red arrow) and a stability loss function (blue arrow). Binding sites are represented as multi-channel 3D images **p***i* and encoded using a CNN *d*_*θ*_ into *n*-dimensional descriptors (filled symbols), which can be compared quickly and easily by computing their pairwise Euclidean distances. The network is trained to make descriptors of matching pocket pairs *P* as similar to each other as possible but to separate the descriptors of non-matching pocket pairs *N* by at least margin distance *m*. In addition, descriptors are encouraged to be robust to small perturbations of the representation, shown as hollow symbols.

### 2.1 Training Dataset

Moving from intuition-based featurization schemes towards learned representations presumes the availability of a large training corpus of pocket pairs with associated 3D structures. To frame the task as a supervised machine learning problem, we assume each pocket pair to be of a certain similarity. Here, we restrict ourselves to the binary case of similar and dissimilar pocket pairs. Unfortunately, obtaining this information is not easy in practice and has been the underlying theme in a range of benchmarking papers. Indeed, the performance of pocket matching algorithms has been shown to depend strongly on the construction of particular datasets (Lee and Im, 2017), and we expect similar behavior to arise when using such datasets for training as well. While highly structurally similar pocket pairs can be easily obtained by considering sequence similar proteins, pairs of unrelated proteins binding similar ligands (Barelier *et al.*, 2015) represent less obvious examples of pocket pairs that may be presumed similar. We expect these cases to represent more closely the needs of desired applications since they are not detectable by methods that rely on sequence-based similarity.

Generally speaking, similarity can be defined on two levels of granularity: for pairs of proteins and for pairs of pockets.

Protein-level similarity is often derived from chemical similarity among the respective binding ligands of two proteins (Keiser *et al.*, 2007) or from commonalities in activity profiles for a set of shared compounds, as measured by functional or binding assays. For example, Chen *et al.* (2016) discriminate protein pairs sharing common active ligands from those without shared active ligands. While the empirical measurement of protein-level similarity is arguably scalable and cost-efficient, pin-pointing the exact binding sites (and protein conformations) responsible for the observed behavior is problematic. This uncertainty makes pocket pair datasets defined at the protein-level unfit for training a pocket-level similarity predictor directly. Nevertheless, such datasets can be used to evaluate pocket matching algorithms by estimating protein similarity as the maximum predicted pocket similarity computed over all pockets and all structures of the respective proteins (Chen *et al.*, 2016).

Pocket-level similarity is derived directly from 3D protein-ligand complexes such as those available in the Protein Data Bank (PDB) (Berman *et al.*, 2000), and is often enhanced by the heuristic assumption that similar ligands bind to similar pockets and vice versa. While providing detailed localization of protein-ligand binding, data acquisition is expensive and pockets are observed in a single bound (*holo-*) conformation, which means that training on such data may not warrant generalization to other induced fit or unbound (*apo-*) protein conformations. Although studies have suggested that conformational rearrangement may be limited (Brylinski and Skolnick, 2008).

Historically, datasets have been constructed as classification experiments involving sets of protein structures bound to a small number of commonly occurring co-crystal ligands (Kahraman *et al.*, 2007; Xie and Bourne, 2008; Hoffmann *et al.*, 2010). However, these datasets tend to be small (hundreds or a few thousand pairs) and may not be sufficiently representative of possible protein binding site space for training purposes. The APoc dataset (Gao and Skolnick, 2013) represents a step towards larger, more general datasets, comprising 34,970 positive and 20,744 negative pairs. Recently, Govindaraj and Brylinski (2018) proposed a large dataset, TOUGH-M1, of roughly one million pairs of protein-ligand binding sites curated from the PDB. Specifically, the authors considered a subset of the PDB including protein structures binding a single “drug-like” ligand. Structures were clustered based on sequence similarity and representative structures bound to a diverse set of ligands were chosen from each cluster. Since the dataset is designed with the prospective use case in mind, in which the location of ligand binding is not available, pocket definition was performed using Fpocket (Le Guilloux *et al.*, 2009) and predicted cavities having the greatest overlap with known binding residues were selected. Finally, bound ligands were also clustered and globally dissimilar protein pairs were identified either within (positive) or between (negative) each ligand cluster. The resulting TOUGH-M1 dataset consists of 505,116 positive pocket pairs and 556,810 negative pocket pocket pairs.

In this work, we choose to train our approach on the TOUGH-M1 dataset. From a machine learning perspective, TOUGH-M1 has the advantages of being large, balanced and offers pocket-level similarities. Notwithstanding, this dataset represents a specific method for defining pocket similarities and it is thus unclear if a trained method can generalize to other datasets, constructed in possibly different ways. We will return to this question in Section 3 and answer it affirmatively.

Finally, let us emphasize that while it is often functional binding sites that are of biological interest, we refer to protein cavities indiscriminately as pockets, since the method discussed is agnostic to the biological relevance of the pockets analyzed.

### 2.2 Volumetric Input Representation

Similarly to recent works addressing pocket detection (Jiménez *et al.*, 2017) and protein-ligand affinity prediction (Jiménez *et al.*, 2018; Stepniewska-Dziubinska *et al.*, 2018), we regard protein structures as 3D images with *c* channels *f* : ℝ^3^ *→* ℝ^*c*^ (4D tensors). This is analogous to the treatment of color images in computer vision as functions assigning a vector of intensities of three primary colors to each pixel, ℝ^2^ *→* ℝ^3^.

There are *c* = 8 feature channels assigned to every point in the 3D image, expressing the presence or absence (occupancy) of atoms in general as well as the presence of atoms exhibiting seven pharmacophoric properties: hydrophobicity, aromaticity, ability to accept or donate a hydrogen bond, positive or negative ionizability, and being metallic. Each atom is thus assigned to at least one feature channel. Occupancy information is given by a smooth indication function of the van der Waals radii of atoms. More precisely, occupancy *f* (**x**)_*h*_ at point **x** *∈* R^3^ in channel *h ∈* {1, .., *c*} corresponds to the strongest indication function over the protein atoms *𝒜*_*h*_ assigned to that channel, formally:

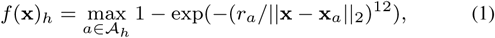

where *r*_*a*_ is van der Waals radius and **x**_*a*_ is the position of atom *a*. Protein structures are retrieved from the PDB, and molecules that are not annotated as part of the main-chain are ignored (*e.g.* water, ligands). This featurization process is analogous to that used for DeepSite (Jiménez *et al.*, 2017) and is based on AutoDock 4 atom types (Morris *et al.*, 2009) and computed using the high-throughput molecular dynamics (HTMD) package (Doerr *et al.*, 2016).

A pocket is represented as tensor **p** *∈* ℝ ^*c×d×d×d*^ created by sampling the corresponding protein structure image *f* over a grid of shape *d × d × d* with step *s*Å. To denote the representation of a particular pocket in *f* centered at point ***µ*** *∈* ℝ^3^ and seen under angle *ϕ*, we use the functional notation **p** = *p*(*f*, ***µ***, *ϕ*). In our datasets of interest, ***µ*** is either the geometric center of a pocket, *i.e.* the centroid of convex hull of alpha spheres in the case of pockets detected with Fpocket 2.0 (Le Guilloux *et al.*, 2009) or the centroid of convex hull of surrounding residues laying within 8 Å of non-hydrogen ligand atoms in the case of pockets defined by their bound ligands.

### 2.3 Learning Pocket Descriptors

We draw inspiration from computer vision, where comparing local image descriptors is the cornerstone of many tasks, such as stereo reconstruction or image retrieval (Szeliski, 2010). There, carefully hand-crafted descriptors such as SIFT (Lowe, 1999) have been recently matched in performance by descriptors learned from raw data (Schönberger *et al.*, 2017; Balntas *et al.*, 2017).

Descriptor learning is usually formulated as a supervised learning problem. Given set 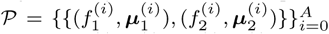 of positive and set 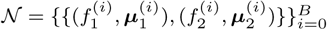 of negative pairs, the goal is to learn a representation such that the descriptors of structurally similar pockets are close to one another in the learned vector space while descriptors of dissimilar pockets are kept far apart. Several objective (loss) functions have been introduced in past work that typically operate on pairs or triplets of descriptors. Triplets are formed by selecting a positive and a negative partner for a chosen anchor (Wang *et al.*, 2014; Hoffer and Ailon, 2015), which is problematic in a pocket matching scenario, as the ground truth relationship between most pocket pairs is unknown: in fact, only 3,991 out of 505,116 positive pairs in TOUGH-M1 can be used for constructing such triplets. Therefore, we build on the pair-wise setup following Simo-Serra *et al.* (2015), which has shown success in computer vision tasks (Balntas *et al.*, 2017). Specifically, given a pair of pockets *Q* = {(*f*_1_, *µ*_1_), (*f*_2_, *µ*_2_)} and orientations *ϕ*_1_, *ϕ*_2_, we minimize the following contrastive loss function (Hadsell *et al.*, 2006) for a pair of pocket representations **p**_1_ = *p*(*f*_1_, ***µ***_1_, *ϕ*_1_) and **p**_2_ = *p*(*f*_2_, ***µ***_2_, *ϕ*_2_):

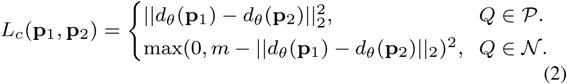

where *d*_*θ*_ : ℝ^*c×d×d×d*^ *→* ℝ^*n*^ is the description function (a neural network) with learnable parameters *θ* computing *n*-dimensional descriptors of pockets. The loss encourages the descriptors of positive pairs to be identical while separating those of negative pairs at least by margin *m >* 0 in Euclidean space. The ability to compute descriptors independently (and in parallel, taking advantage of modern GPUs) and compare them efficiently by evaluating the L2 norm of their difference is very advantageous especially for large-scale searches and all-against-all scenarios.

### 2.4 Towards Descriptor Robustness

A highly desirable property of pocket matching tools is robustness with respect to the the chosen pocket representation and the inherent variability in pocket definition. In particular, this includes discretization artifacts due to input grid resolution *s*, the orientation of pockets in 3D space *ϕ* (there is no canonical orientation of proteins nor their pockets in space) and their precise delimitation (the inclusion or exclusion of a small number of atoms) affecting the position of their geometric centers ***µ***. Robustness here would also render the network stable to protein conformational variability.

Robustness has been traditionally addressed by using fuzzy featurization schemes and explicit alignment techniques in previous pocket matching tools, see Naderi *et al.* (2018) for a review, and by using data augmentation in machine learning methods (Kauderer-Abrams, 2017). The latter strategy is also applicable in our case, where data augmentation amounts to randomly sampling *ϕ* (implemented as random rotation around a random axis) and adding a random vector *ϵ, ∥ ϵ ∥*_2_ *≤* 2 Å, to ***µ*** for each pocket seen during training in order to stimulate the descriptor function *d*_*θ*_ to become invariant to such perturbations. However, we have not been able to achieve a sufficient level of invariance in practical experiments with only this approach, which we consider related to series of observations of the vulnerability of neural networks to small geometric input transformations in both adversarial (Fawzi and Frossard, 2015) and benign settings (Azulay and Weiss, 2018).

This motivates us to introduce an additional, explicit stability objective (Zheng *et al.*, 2016). Given two perturbed representations of the same pocket (*f*, ***µ***), 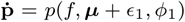 and 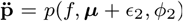, we encourage their descriptors to be identical by minimizing the following stability loss:

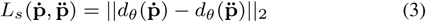

The contrastive and the stability loss are then minimized jointly in a linear combination weighted with hyperparameter *λ >* 0 as:

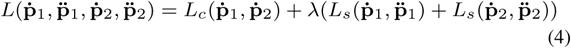

### 2.5 Network Architecture

Our description function *d*_*θ*_ is a relatively shallow CNN. CNNs are hierarchical machine learning models consisting of layers of several types, see *e.g.* Goodfellow *et al.* (2016) for an overview. To support the above mentioned desire for translationally and rotationally invariant descriptors, we draw on the recent progress in learning rotationally equivariant features. Concretely, we use 3D steerable CNNs (Weiler *et al.*, 2018), where 3D convolutional filters are parameterized as a linear combination of a complete steerable kernel basis. Such a technique for parameter sharing allowed us to considerably decrease the number of learnable parameters down to the order of 105 and therefore to reduce possible overfitting.

The network, described in detail in Table 1, consists of six convolutional layers. We prefer striding preceded by low-pass filtering, as recommended by Azulay and Weiss (2018), over pooling, which has empirically led to more stable networks. The computed descriptors are additionally normalized to have unit length, as *per* usual practice (Lowe, 1999; Balntas *et al.*, 2017).

**Table 1.** The architecture of network used in the experiments, in top down order. SCB denotes a steerable 3D convolution block with batch normalization and ReLU scalar and sigmoid gate activation (Weiler et al., 2018) and C denotes a standard 3D convolution layer preceded by ReLU activation and batch normalization (Ioffe and Szegedy, 2015).

|  |  |
| --- | --- |
| SCB | kernel size 7, padding 3, stride 2, $4 \times 16$ fields of order 0-3 |
| SCB | kernel size 3, padding 1, stride 1, $4 \times 32$ fields of order 0-3 |
| SCB | kernel size 3, padding 1, stride 2, $4 \times 48$ fields of order 0-3 |
| SCB | kernel size 3, padding 0, stride 1, $4 \times 64$ fields of order 0-3 |
| SCB | kernel size 3, padding 0, stride 2, 256 fields of order 0 |
| C | kernel size 1, padding 0, stride 1, $n$ output channels |

### 2.6 Training Details

Besides the strategies for rotational and translational data augmentation described above, random points are sampled with probability 0.1 instead of pocket centers ***µ*** in negative pairs to increase the variability of negatives and regularize the behavior of the network outside pockets over the whole protein. We set margin *m* = 1, loss weight *λ* = 1 and descriptor dimensionality *n* = 128. Networks are trained on balanced batches of 16 quadruples for 6000 iterations with a variant of stochastic gradient descent, Adam (Kingma and Ba, 2014), with weight decay of 5 *×* 10^−4^ and learning rate of 0.001 step-wise annealed after 4000 iterations. Training takes about 1.5 days on a single GPU. We observe that higher resolution and larger spatial context are generally beneficial and set *d* = 24 and *s* = 1 Å as a compromise between computational efficiency and performance in this work. Finally, let us remark that we use the same architecture and training parameters for all networks presented in this work.

## 3. Results and Discussion

### 3.1 TOUGH-M1 Dataset

TOUGH-M1 (Govindaraj and Brylinski, 2018) is a dataset of 505,116 positive and 556,810 negative protein pocket pairs defined from 7,524 protein structures. Pockets are defined computationally with Fpocket 2.0 (Le Guilloux *et al.*, 2009) and filtered to include only predicted cavities having the greatest overlap with known binding residues, see Section 2.1. As the TOUGH-M1 dataset is used for both training and evaluation but has not been used in a machine learning setting previously, we first define a sensible data splitting strategy before we report our results and compare to several baseline methods.

#### Training and evaluation strategy

The definition of independent subsets of data, as desired for meaningful evaluation of machine learning-based methods, is not straightforward in this case. Indeed, a number of recent works have commented on the need for robust splitting methodologies when working with protein structure data (Li and Yang, 2017; Feinberg *et al.*, 2018). A single protein structure may take part in multiple pairwise relationships, some possibly being in the training set and some in the test set, leading to a potential for information leakage. In addition, TOUGH-M1 may contain multiple structures representing a single protein family. Here, we propose to split at the structure level (instead of pairs), devoting 80% of structures for training and hyperparameter tuning and reserving the remaining 20% (1,457 structures, 40,296 pairs) as a hold-out test set. Any pairs connecting training and testing protein structures are discarded. Structures assigned to a common UniProt accession number (determined using the SIFTS database (Velankar *et al.*, 2012)) are always allocated to the same subset. 220 PDB IDs to which no UniProt accession number could be assigned were removed.

After hyperparameter search rounds, we retrain our network on the whole training set (5,847 structures, 628,234 pairs). Following Govindaraj and Brylinski (2018), the performance is measured with the Receiver Operating Characteristic (ROC) and the corresponding Area Under the Curve (AUC) is reported. To estimate the sensitivity to the choice of particular splits, we use repeated random sub-sampling (Monte Carlo) validation (Dubitzky *et al.*, 2007). Having finalized the hyperparameters, we repeat splitting and training on ten random permutations of TOUGH-M1 and measure the standard error over the respective test sets.

#### Baselines

We compare DeeplyTough to three alignment-based methods chosen by the authors of the TOUGH-M1 dataset and to an additional alignment-free approach. *APoc* (Gao and Skolnick, 2013) optimizes the alignment between pocket pairs by iterative dynamic and integer programming, considering the secondary structure and fragment fitting. *G-LoSA* (Lee and Im, 2017) uses iterative maximum clique search and fragment superposition. *SiteEngine* (Shulman-Peleg *et al.*, 2005) uses geometric hashing and matching of triangles of physicochemical property centers. Last, *PocketMatch* (Yeturu and Chandra, 2008) is an alignment-free method which represents pockets as lists of sorted distances encoding their shape and chemical properties. While we reuse the list of matching scores published by Govindaraj and Brylinski (2018) for each alignment-based method and split them into folds according to our evaluation strategy, we compute matching scores for PocketMatch ourselves.

#### Results

The measured ROC curves for the TOUGH-M1 dataset are shown in Figure 2. DeeplyTough achieves an AUC of 0.933, outperforming by far all other approaches and achieving a substantial improvement over the second best performing method, SiteEngine (AUC 0.733). The remaining two alignment-based methods, G-LoSA and APoc, achieve greater AUCs than PocketMatch, the alignment-free approach, with AUCs of 0.683, 0.662 and 0.611, respectively. Furthermore, the performance of all methods is fairly stable across different test (and training) sets, with DeeplyTough achieving the lowest standard error.

**Fig. 2.**
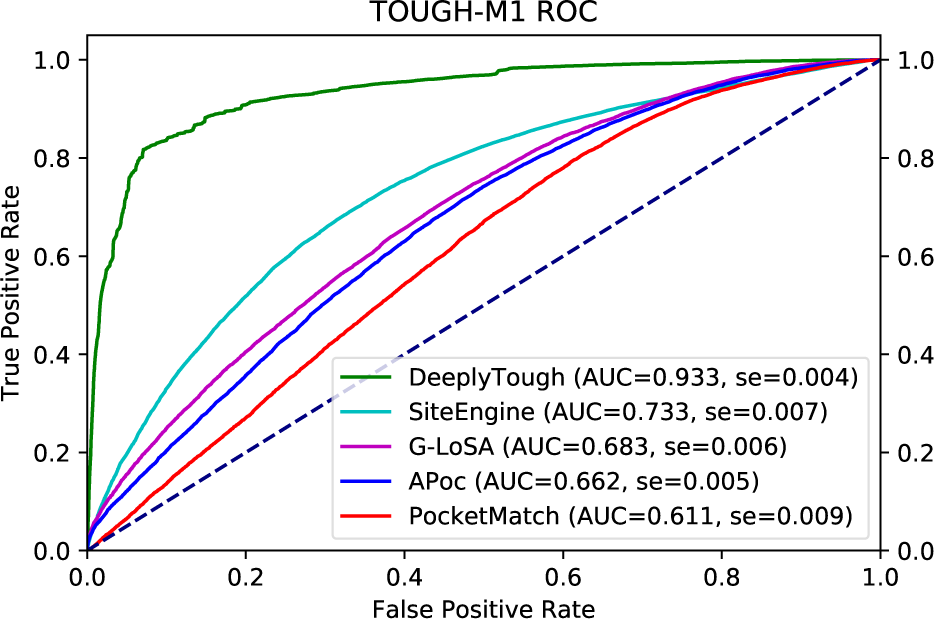
ROC plot with associated AUC values evaluating the performance of pocket matching algorithms on the TOUGH-M1 test set (40,296 pairs). Standard error, denoted as se, is measured over ten random splits. The dashed line represents random predictions.

Analysis of the TOUGH-M1 positive pocket pairs that were assigned large distances (false negatives) highlights potentially ambiguous ground truths in the dataset. In particular, the bound ligands of false negative pockets show enrichment of biologically versatile endogenous molecules such as nucleotides (ATP, ACO), amino acid monomers (TYR, PRO, ASP) and sugars (GLC, NDG), as well as a number of non-biologically relevant ligands involved in the production of protein structure data such as crystallization agents (MPD, MRD), purification agents (CIT, FLC) and buffer solutions (TRS, B3P). While these pocket pairs do represent instances where related ligands are bound to unrelated proteins (constituting the definition of a positive pocket pair), we argue that in some cases there is limited structural similarity between pockets, and shared binding may be attributed to the conformational flexibility (Haupt *et al.*, 2013) or non-specificity of the bound ligand. These cases may be considered a limitation of the current dataset. On the other hand, analysis of TOUGH-M1 negative pocket pairs that were assigned small distances (false positives), revealed a large interconnected network of pocket pairs that were predicted similar. Inherently, there is a high potential for false positives in pocket pair datasets since the absence of common bound ligands in extant data does not render it impossible for two binding sites to bind related ligands. Of the false positive pocket pairs examined, we found that many pockets were bound to polar ligand moieties containing anionic groups such as phosphates (2P0, T3P, S6P), sulfonamides (E49, 3JW), and carboxylates (G39, BES). Furthermore, false positive pockets seem to be enriched with polar residues suggesting that there may be similarity between these pockets, despite their negative labels.

DeeplyTough performs well on the TOUGH-M1 dataset, however, it must be remembered that this is a trained method, and while results are calculated for data held-out from training, the data is still drawn from the same underlying distribution. It is therefore interesting to evaluate the performance of DeeplyTough on further datasets constructed independently.

### 3.2 Vertex Dataset

The Vertex dataset introduced by Chen *et al.* (2016) comprises 6,598 positive and 379 negative protein pairs defined from 6,029 protein structures. The protocol for annotation of protein pairs derives from commonalities (or lack thereof) among experimentally measured ligand activities. In this benchmark, predicted protein-level similarities are obtained from a set of pocket-level similarities. In particular, the smallest of *k × l* predicted pocket distances is assigned to each protein pair of interest with *k* ligand binding sites collected across different PDB structures for one protein and *l* for the other. For the Vertex dataset, this amounts to 1,461,668 positive and 102,935 negative pocket comparisons in total.

Unlike in TOUGH-M1, where binding sites are obtained from predicted cavities, the Vertex dataset defines pockets using their bound ligands directly. Specifically, we define a pocket as all complete protein residues with any atom falling within 8 Å of any ligand atom.

#### Training and evaluation strategy

The network is trained on the whole TOUGH-M1 dataset. However, to prevent information leakage, we discard all structures having their UniProt accession number found in the Vertex dataset as well as 220 PDBs to which no UniProt accession number could be assigned, resulting in 6,979 structures and 820,140 pairs left for training. Following Chen *et al.* (2016), we measure performance with the ROC curve and corresponding AUC.

#### Baselines

We compare our approach to *SiteHopper* (Batista *et al.*, 2014), a structure-based pocket matching method chosen by the authors of the dataset. SiteHopper is an alignment-based method which represents binding sites as sets of points describing the molecular surface and nearby physicochemical features, which are aligned by maximizing the overlap of point-centered Gaussian functions. We also compare to PocketMatch, as for the TOUGH-M1 analysis. G-LoSA was omitted from this study due to running time of hundreds of days on a single processor. Results for SiteHopper were kindly provided in personal communication by Chen *et al.* (2016).

#### Results

The measured ROC curves are shown for the Vertex dataset in Figure 3. We can see that both SiteHopper (AUC 0.887) and DeeplyTough (AUC 0.818) achieve good performance on the dataset, while the gap to PocketMatch is large (AUC 0.604). Importantly, the result indicates that our method generalizes well across two different methods for defining binding site geometric centers (computational and ligand-based). These results also hint that the ground truth pocket similarities in TOUGH-M1 and Vertex are comparable, despite being derived by different protocols.

**Fig. 3.**
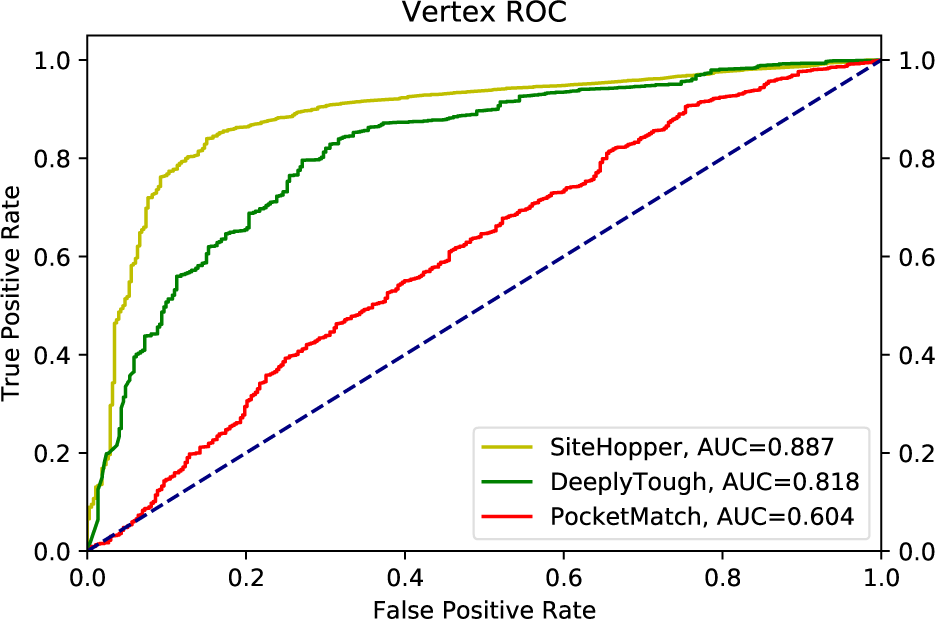
ROC plot with associated AUC values evaluating the performance of pocket matching algorithms on the Vertex dataset (6,977 protein pairs). The dashed line represents random predictions.

### 3.3 ProSPECCTs Datasets

ProSPECCTs (Ehrt *et al.*, 2018) is a collection of ten benchmarks recently assembled to better understand the performance of pocket matching for various practical applications. As for the Vertex dataset above, binding sites are defined by their bound ligands and we include complete protein residues with any atom that falls within 8 Å of any ligand atom.

#### Training and evaluation strategy

The network is trained on the whole TOUGH-M1 dataset. To prevent information leakage, we discard all structures having their UniProt accession number found in any of the ProSPECCTs datasets as well as 220 PDB IDs to which no UniProt accession number could be assigned, resulting in 6,369 structures and 715,520 pairs left for training. Following Ehrt *et al.* (2018), the performance is measured with ROC curves and the corresponding AUC.

#### Baselines

We compare our approach to a diverse set of 21 pocket matching methods chosen by the authors of the benchmark, directly reusing their published results.

#### Results

The measured AUC scores for the ProSPECCTs datasets are given in Table 2 and compactly visualized in Figure 4.

**Table 2.** AUC values for 22 pocket matching methods on each of ten ProSPECCTs datasets.

|  | P1 | P1.2 | P2 | P3 | P4 | P5 | P5.2 | P6 | P6.2 | P7 |
| --- | --- | --- | --- | --- | --- | --- | --- | --- | --- | --- |
| Cavbase | 0.98 | 0.91 | 0.87 | 0.65 | 0.64 | 0.60 | 0.57 | 0.55 | 0.55 | 0.82 |
| FuzCav | 0.94 | 0.99 | 0.99 | 0.69 | 0.58 | 0.55 | 0.54 | 0.67 | 0.73 | 0.77 |
| FuzCav (PDB) | 0.94 | 0.99 | 0.98 | 0.69 | 0.58 | 0.56 | 0.54 | 0.65 | 0.72 | 0.77 |
| Grim | 0.69 | 0.97 | 0.92 | 0.55 | 0.56 | 0.69 | 0.61 | 0.45 | 0.65 | 0.70 |
| Grim (PDB) | 0.62 | 0.83 | 0.85 | 0.57 | 0.56 | 0.61 | 0.58 | 0.45 | 0.50 | 0.64 |
| IsoMIF | 0.77 | 0.97 | 0.70 | 0.59 | 0.59 | 0.75 | 0.81 | 0.62 | 0.62 | 0.87 |
| KRIPO | 0.91 | 1.00 | 0.96 | 0.60 | 0.61 | 0.76 | 0.77 | 0.73 | 0.74 | 0.85 |
| PocketMatch | 0.82 | 0.98 | 0.96 | 0.59 | 0.57 | 0.66 | 0.60 | 0.51 | 0.51 | 0.82 |
| ProBiS | 1.00 | 1.00 | 1.00 | 0.47 | 0.46 | 0.54 | 0.55 | 0.50 | 0.50 | 0.85 |
| RAPMAD | 0.85 | 0.83 | 0.82 | 0.61 | 0.63 | 0.55 | 0.52 | 0.60 | 0.60 | 0.74 |
| Shaper | 0.96 | 0.93 | 0.93 | 0.71 | 0.76 | 0.65 | 0.65 | 0.54 | 0.65 | 0.75 |
| Shaper (PDB) | 0.96 | 0.93 | 0.93 | 0.71 | 0.76 | 0.66 | 0.64 | 0.54 | 0.65 | 0.75 |
| VolSite/Shaper | 0.93 | 0.99 | 0.78 | 0.68 | 0.76 | 0.56 | 0.58 | 0.71 | 0.76 | 0.77 |
| VolSite/Shaper (PDB) | 0.94 | 1.00 | 0.76 | 0.68 | 0.76 | 0.57 | 0.56 | 0.50 | 0.57 | 0.72 |
| SiteAlign | 0.97 | 1.00 | 1.00 | 0.85 | 0.80 | 0.59 | 0.57 | 0.44 | 0.56 | 0.87 |
| SiteEngine | 0.96 | 1.00 | 1.00 | 0.82 | 0.79 | 0.64 | 0.57 | 0.55 | 0.55 | 0.86 |
| SiteHopper | 0.98 | 0.94 | 1.00 | 0.75 | 0.75 | 0.72 | 0.81 | 0.56 | 0.54 | 0.77 |
| SMAP | 1.00 | 1.00 | 1.00 | 0.76 | 0.65 | 0.62 | 0.54 | 0.68 | 0.68 | 0.86 |
| TIFP | 0.66 | 0.90 | 0.91 | 0.66 | 0.66 | 0.71 | 0.63 | 0.55 | 0.60 | 0.71 |
| TIFP (PDB) | 0.55 | 0.74 | 0.78 | 0.56 | 0.57 | 0.54 | 0.53 | 0.56 | 0.61 | 0.66 |
| TM-align | 1.00 | 1.00 | 1.00 | 0.49 | 0.49 | 0.66 | 0.62 | 0.59 | 0.59 | 0.88 |
| DeeplyTough | 0.96 | 0.97 | 0.93 | 0.72 | 0.72 | 0.66 | 0.61 | 0.59 | 0.59 | 0.80 |
| rank | 7-10 | 12-14 | 11-13 | 5 | 8 | 6-9 | 8-9 | 8-9 | 13-14 | 8-9 |

P4, respectively), consistent with the intuition that pockets that have been more heavily mutated should be easier to differentiate.

**Fig. 4.**
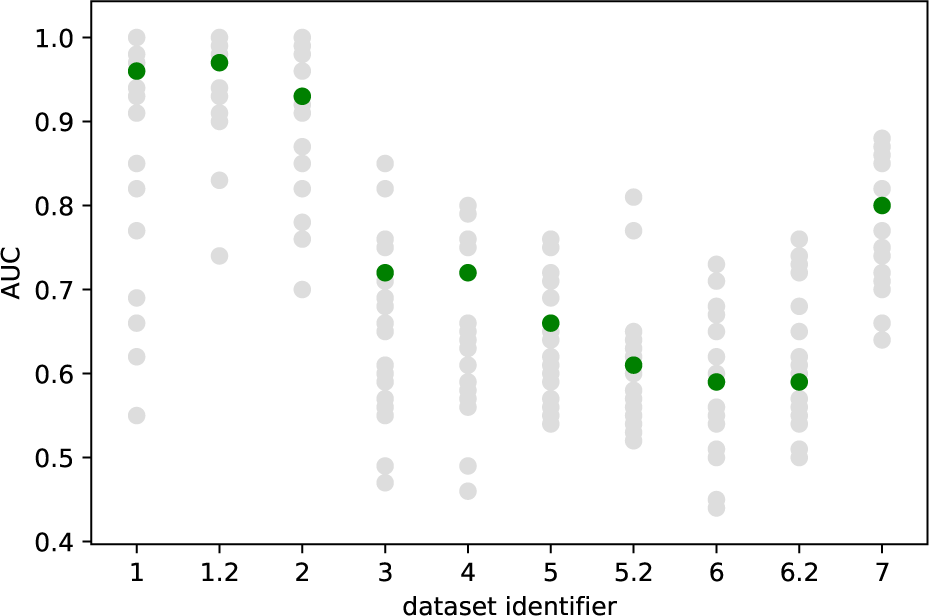
AUC values for DeeplyTough (green) and 21 other pocket matching methods (gray) on each of ten ProSPECCTs datasets.

Dataset P1 evaluates the sensitivity to the binding site definition by comparing structures with identical sequences which bind to chemically different ligands at identical sites. Dataset P1.2 measures this exclusively for chemically similar ligands. By reaching AUCs of 0.96 and 0.97, respectively, we can conclude that DeeplyTough is fairly robust to varying pocket definitions, which may be attributed to our stability loss as well as our data augmentation strategy. In Table 3, we observe that the stability loss alone is responsible for more than AUC 0.1 difference across multiple ProSPECCTs datasets. In addition, the box plot in Figure 5 illustrates a clear distance-dependent distinction between identical and non-matching binding site pairs, likely a virtue of our margin-based contrastive loss function.

**Table 3.**
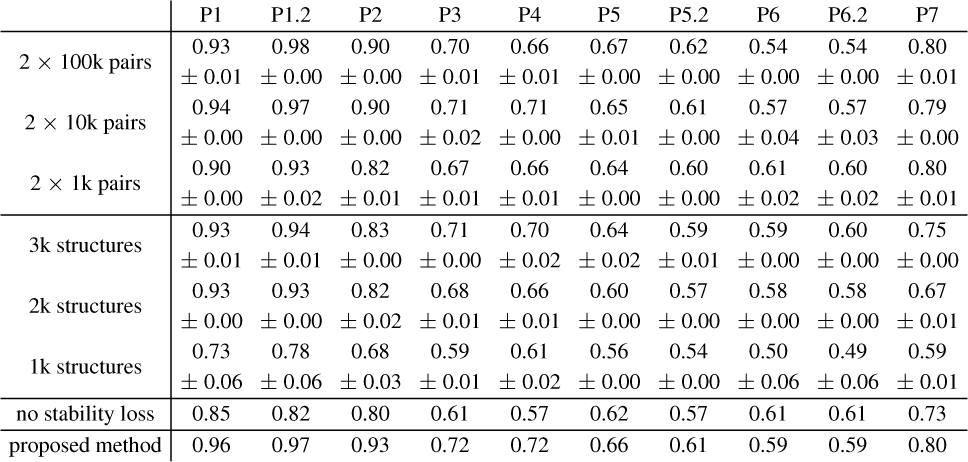
The effect of training dataset size, expressed by the amount of positive and negative binding site pairs as well as unique PDB structures, and of stability loss measured in AUC values on each of ten ProSPECCTs datasets, with standard error if sensible.

**Fig. 5.**
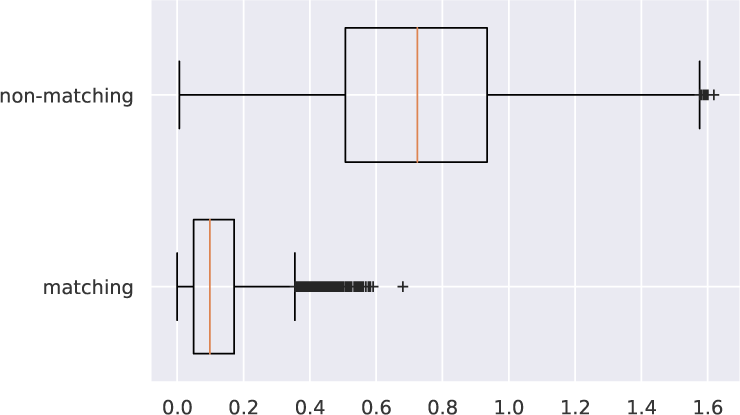
The distribution of distances between DeeplyTough descriptors of matching and non-matching binding sites of structures with identical sequences (ProSPECCTs Dataset P1).

Dataset P2 assesses the sensitivity to binding site flexibility by comparing the pockets of Nuclear Magnetic Resonance (NMR) structures with more than one model in the structure ensemble. DeeplyTough achieves AUC 0.93, which indicates a slight susceptibility to the conformational variability of proteins. We believe this could be addressed by introducing an appropriate data augmentation strategy in the training process.

Next, two decoy datasets evaluate the discrimination between nearly identical binding sites differing by five artificial mutations leading to different physicochemical properties (Dataset P3) or both physicochemical and shape properties (Dataset P4). Performing at AUC 0.72, DeeplyTough has some difficulty ranking original binding sites pairs with identical sequences, higher than pairs consisting of an original structure and a decoy structure. This suggests that the learned network might be overly robust and may not pay enough attention to modifications in the sites. When compared to existing approaches, however, our approach ranks proportionally well as fifth and eighth, respectively. Moreover, the performance is well correlated with the number of mutations in the sites (AUC 0.55, 0.63, 0.65, 0.72 and AUC 0.53, 0.60, 0.66, 0.68 for one to four mutations in Dataset P3 and P4, respectively), consistent with the intuition that pockets that have been more heavily mutated should be easier to differentiate.

A further two datasets contain sets of dissimilar proteins binding to identical ligands and cofactors. Datasets P5 and P5.2 have been compiled by Kahraman *et al.* (2007) and contain 100 structures bound to one of ten ligands (excluding and including phosphate binding sites, respectively). Datasets P6 and P6.2 contain pairs of unrelated proteins bound to identical ligands, assembled by Barelier *et al.* (2015) (excluding and including cofactors, respectively). Our method scores better on the kahraman dataset (AUC 0.66) than on unrelated proteins (AUC 0.59), consistent with reports that the kahraman dataset represents an easy benchmark since the chosen ligands may be distinguished solely by their sizes (Hoffmann *et al.*, 2010).

Finally, Dataset P7 is a retrieval experiment measuring the recovery of known binding site similarities within a set of diverse proteins. DeeplyTough reaches AUC 0.80 (average precision 0.44), which places among the best performing half of the baseline approaches.

In summary, our method performs consistently well across ProSPECCTs datasets although it does not attain the best performance for any individual task. For practical applications, we suggest these results support the use of DeeplyTough as a fast basic universal tool, rather than a specialist one.

#### Running time

In addition, Ehrt *et al.* (2018) published the running times for each algorithm on Dataset P5 (100 pockets and 10,000 comparisons). The runtime for DeeplyTough is 206.4 seconds in total, where the preprocessing with AutoDock 4 (Morris *et al.*, 2009) and HTMD (Doerr *et al.*, 2016) requires 191.4 seconds (serialized on a single CPU core) and the descriptor computation and comparison takes 15 seconds on an Nvidia Titan X. This makes ours the fourth fastest approach in the benchmark, behind PocketMatch, RAPMAD, and TM-align. SiteHopper is a slower approach with a total runtime of 3982.6 seconds (17th of the ProSPECCTs baseline methods). For DeeplyTough, we expect that even further reduced runtimes may be achieved through full parallelism of the initial preprocessing.

#### Protein binding site space

T-distributed Stochastic Neighbor Embedding (t-SNE) (Maaten and Hinton, 2008) can be used to visualize the learned descriptor space obtained by DeeplyTough. Figure 6 shows the embeddings of pockets in Dataset P1 colored by their UniProt accession numbers. For the most part, protein pockets derived from the same protein family are clustered together, suggesting that the network embeds similar pockets close to each other in the descriptor space. Similar conclusions can be drawn for protein pockets derived from the Vertex dataset in Figure 7, wherein pockets are colored by their respective top-level SCOPe classifications (Fox *et al.*, 2013). These embeddings suggest that DeeplyTough can be useful for large-scale analyses of protein binding site space (Meyers *et al.*, 2016).

**Fig. 6.**
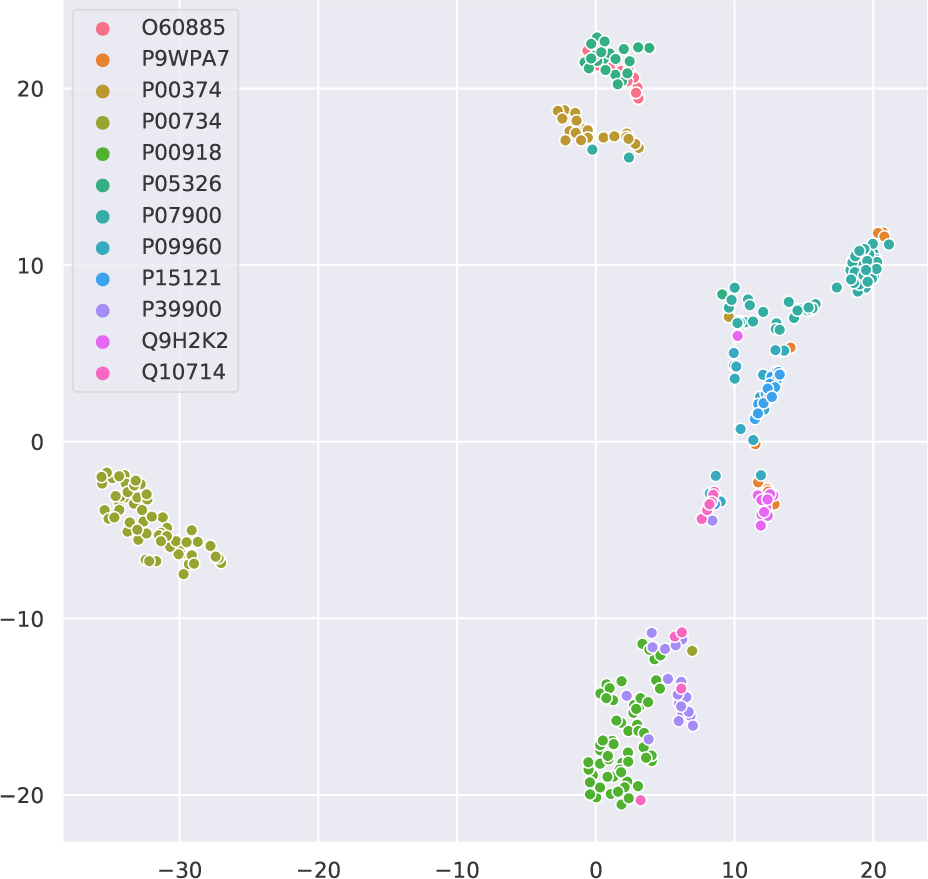
2D t-SNE visualization of descriptors of binding sites in 12 protein groups, denoted by their UniProt accession number, comprised in ProSPECCTs Dataset P1.

**Fig. 7.**
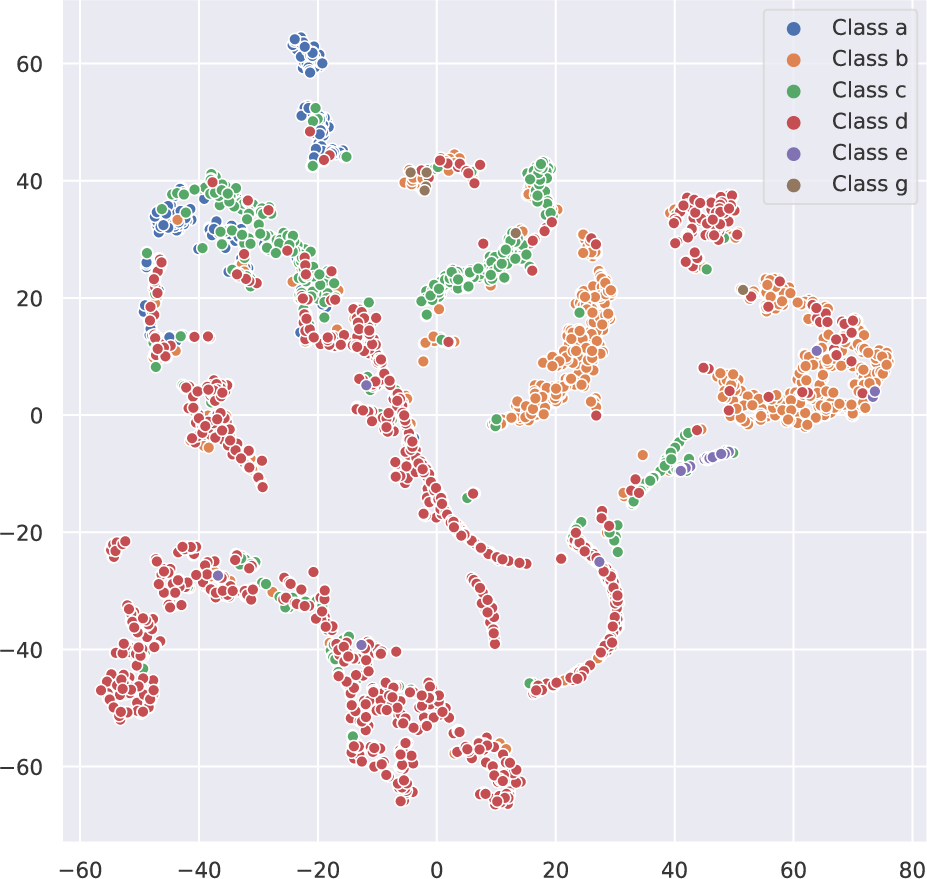
2D t-SNE visualization of descriptors of binding sites in the Vertex dataset, labeled by the top-level SCOPe class of their proteins.

### 3.4 Training Data Ablation

Deep learning models are proverbially known to require large amounts of data for training. To provide an insight into the dependence of DeeplyTough on the large scale of the TOUGH-M1 dataset, we experiment by artificially limiting the training dataset size using two approaches, in both cases validating on ProSPECCTs as an independent set. For simplicity, all training and network hyperparameters are kept fixed.

First, we investigate restricting the number of relationships the network is allowed to see. Thus, pairs of random subsets of size between 1,000 and 100,000 are sampled from the positive and negative set of pocket pairs present in the original training set. The results shown in the upper part of Table 3 suggest the network does not strongly suffer from the removal of training data in this way, even if the training set is smaller by two orders of magnitude. We expect the likely cause of this is that even for a reduced training set of only 2,000 pairs, the effective number of structures is still relatively high (about 2,200 PDBs).

Hence, we look into constraining the pocket variability in the data. Random subsets of size varying between 1,000 and 3,000 are sampled from the original 6,369 structures available for training and only pairs lying within these subsets are retained. The results shown in the bottom part of Table 3 indicate that the performance starts to severely deteriorate once the number of structures drops below 2,000, even if this corresponds to about 70,000 induced pairs. Therefore, we may conclude that for our method, pocket diversity in the data is relatively more important than the number of ground truth relationships. This observation suggests that it may be appropriate to construct new pocket matching datasets using as many structures from the PDB as possible, even if relatively few pocket pairs are defined.

Finally, we reverse the settings and train on ProSPECCTs while validating on the TOUGH-M1 and Vertex datasets (training on 77,665 pairs over 1,395 structures for the former and on 30,175 pairs over 1,246 structures for the latter). This results in major degradation of AUC scores, 0.740 on TOUGH-M1 (*versus* 0.933 when training on TOUGH-M1) and 0.709 on the Vertex dataset (*versus* 0.817). To summarize, it appears that the large scale of TOUGH-M1 is a necessary condition for our method to perform well. However, we expect pre-trained DeeplyTough to be amenable to successful fine-tuning for smaller task-specific datasets.

## 4 Conclusion

In this work we have proposed a deep learning-based approach for pocket matching. DeeplyTough encodes the 3D structure of protein binding sites using a CNN into descriptors such that the similarity between binding sites is reflected in the Euclidean distances between their descriptors. Once a set of descriptors is computed, pocket matching is simple and efficient, without any alignment operation taking place. In a thorough evaluation on three benchmark datasets, we have demonstrated excellent performance on held-out data coming from the training distribution (TOUGH-M1) and competitive performance when the trained network needs to generalize to independently constructed datasets (Vertex, ProSPECCTs). We have taken advantage of several recent innovations such as rotationally and translationally invariant CNNs, data augmentation and the inclusion of an explicit stability loss function to encourage robustness of the network with respect to nuisance variability of the input. Overall, we expect trained methods for pocket matching to remove the bias associated with intuition-based featurization schemes, and also enable effective large scale binding site analyses.

Having presented one of the first trainable methods for pocket matching, there are many exciting avenues of future research. Exploring different methods for obtaining supervision is perhaps the most promising direction. For example, binary labels could be replaced with continuous labels based on chemical similarity of ligands. In addition, the problem could be cast as multiple-instance learning in order to use protein-level similarity coming from assays as a form of weak supervision. Another direction is to investigate other input representations, such as graphs or surfaces. Finally, experiments with various model explainability techniques will be important for giving practitioners insights into the currently rather black-box nature of the algorithm.

## Acknowledgments

We thank Marwin Segler, Mohamed Ahmed, Nathan Brown and Amir Saffari for helpful discussions, as well as Lidio Meireles for providing us with results for SiteHopper.

